# OrganoLabeling: Quick and accurate annotation tool for organoid images

**DOI:** 10.1101/2024.04.16.589852

**Authors:** Burak Kahveci, Elifsu Polatli, Yalin Bastanlar, Sinan Guven

**Affiliations:** Izmir Biomedicine and Genome Center, Izmir, Türkiye; Izmir International Biomedicine and Genome Institute, Dokuz Eylul University, Izmir, Türkiye; Department of Computer Engineering, Izmir Institute of Technology, Izmir, Türkiye; Faculty of Medicine, Medical Biology and Genetics Department, Dokuz Eylul University, Izmir, Türkiye

**Keywords:** organoid, embryoid body, image processing, deep learning

## Abstract

Organoids are self-assembled 3D cellular structures that resemble organs structurally and functionally, providing *in vitro* platforms for molecular and therapeutic studies. Generation of organoids from human cells often require long and costly procedures with arguably low efficiency. Prediction and selection of cellular aggregates that result in healthy and functional organoids can be achieved using artificial intelligence-based tools. Transforming images of 3D cellular constructs into digitally processible datasets for training deep learning models require labeling of morphological boundaries, which often is performed manually. Here we report an application named OrganoLabeling, which can create large image-based datasets in consistent, reliable, fast, and user-friendly manner. OrganoLabeling can create segmented versions of images with combinations of contrast adjusting, K-means clustering, CLAHE, binary and Otsu thresholding methods. We created embryoid body and brain organoid datasets, of which segmented images were manually created by human researchers and compared with OrganoLabeling. Validation is performed by training U-Net models, which are deep learning models specialized in image segmentation. U-Net models, that are trained with images segmented by OrganoLabeling, achieved similar or better segmentation accuracies than the ones trained with manually labeled reference images. OrganoLabeling can replace manual labeling, providing faster and more accurate results for organoid research free of charge.

**Translational Impact:** We developed an image processing-based tool called OrganoLabeling generating datasets to train deep learning models and achieved its performance by comparing with experienced researchers. Here we demonstrate and validate OrganoLabeling, a tool that is as fast and successful as humans, automating the process of creating datasets for use in training deep learning models that can be used for disease analysis and translational purposes in medicine. OrganoLabeling can be broadly applied in artificial intelligence engaged life sciences focusing on stem cell based organoid research.

**Graphical Abstract:** 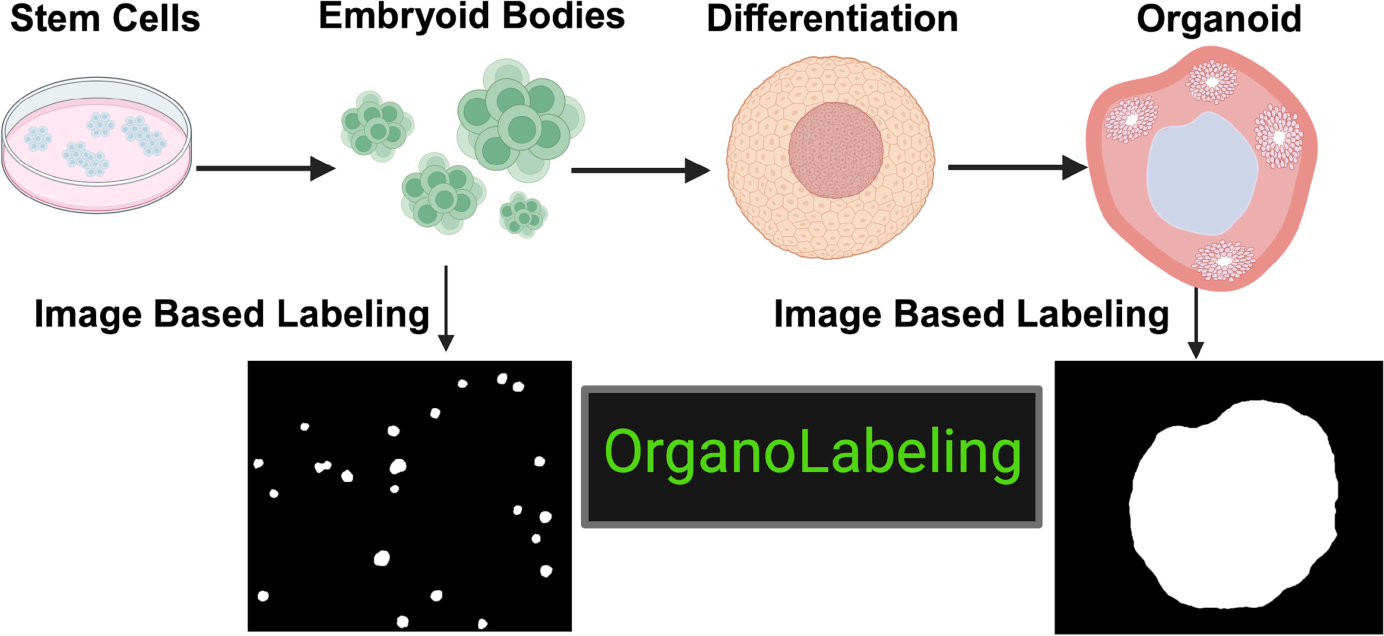

## 1. Introduction

Organoids are 3D tissue equivalents that provide the structural and functional recapitulation of an organ that can be generated from stromal or pluripotent stem cells [1]. Organoids provide *in vitro* venues to investigate molecular and mechanistic studies on target organ including realistic models for diseases and drug testing. Advances in stem cell research, bioengineering and organoid technologies lead the generation of diverse portfolio of organoids including brain, intestine, liver, kidney, stomach, lung, thyroid and lacrimal gland [1–11]. Brain organoids (BOs) resemble embryonic development of the cerebellum and can be employed in understanding neurodegenerative diseases and brain research [12].

Generation of BOs morphologically starts with self-aggregation of cells followed by formation of embryoid bodies (EB) that hold capacity to differentiate to three distinct germ layers. Neural induction of EBs followed with further differentiation and maturation leads the formation of BOs. Time to the generate human organoids and costs associated with the process urges the experimental planning and data process to be well optimized minimizing human error [13].

Machine learning (ML) and deep learning (DL), sub-fields of artificial intelligence, can be used in every field from health to finance, from education to security and increase the quality of life [14]. Traditional ML methods have disadvantages such as requiring feature extraction. Deep learning methods provide automatic feature extraction and capacity to handle operations demanding high processing power such as image data [15, 16]. In recent years, DL models have been used frequently for segmentation tasks [17, 18]. U-Net [19] is a DL architecture that has gained significant popularity in medical image segmentation. However, the segmentation performance of these DL models depends on the segmentation quality of the images in the training sets. The performance of these models may be limited due to biases that may occur in images manually tagged by humans [20]. In biological sciences, images to be used for segmentation tasks can be noisy due to the complexity of the microenvironment of tissues or cells imaged, making labeling by humans difficult or biased [21]. Moreover, since DL models need to be trained with large datasets in order to best adapt to real-life scenarios, the process of labeling these datasets is quite time-consuming [22]. Automatization of the labeling process becomes essential for eliminating human bias and accelerating the process.

A limited number of AI studies have been conducted using organoid images in the literature [23–28]. Additionally, studies such as OrganoID[29] and OrgaExtractor[30] have been developed to perform morphology analysis using deep learning models (Table 1).

**Table 1.** Image processing studies for organoid image segmentation.

| Dataset | Goal | Method | References |
| --- | --- | --- | --- |
| Brain Organoids | Segmentation | K-means, SVM | [23] |
| Brain Organoids | Segmentation | U-Net | [24] |
| Neurons and human mammary gland acinar organoids | Clustering & Classification | Watershed Algorithm | [25] |
| Bladder cancer organoids | Image Processing - Texture Based Analysis & Classification | Image features similarity | [26] |
| Bladder cancer organoids | Segmentation | U-Net | [27] |
| Mouse-Organoid-Cells | Segmentation | ERF-Net | [28] |
| Colon Organoids | Segmentation | U-Net | [30] |
| Pancreatic<br>Cancer and<br>colon<br>Organoids | Segmentation | U-Net | [29] |

In this study, we developed OrganoLabeling, robust, publicly available and no-code labeling tool based on image processing. OrganoLabeling accelerates the process of generating bias-free ground truth images to be employed by DL models. Its user-adjustable parameters make OrganoLabeling easy to adapt in training set creation tasks for analysis of organoid and medical images. We have validated increased performance of segmented images created with OrganoLabeling compared to reference manually segmented images.

## 2. Methods

### 2.1 Generation of brain organoids from human induced pluripotent stem cells (hiPSC)

Human induced pluripotent stem cells (hiPSCs) obtained from healthy donors with informed consent are kindly provided from Prof. Tamer Onder [31]. hiPSC aggregates were plated on 1% Matrigel hESC-Qualified Matrix (Corning, Cat no. 354277) coated plates and cultured in mTeSR1 (STEMCELL Technologies, Cat no. 85850) with daily medium change. Morphologically differentiated cells were scratched out with pipet tip. hiPSCs were passaged upon 80% confluency with ReLeSR (STEMCELL Technologies, Cat no. 05872).

For the collection of EBs datasets embryoid bodies were generated according to protocol based on [32]. First, hiPSCs were plated as cell clumps on low attachment plates and cultured with embryonic stem cell medium (ESCM) composed of DMEM/F-12 (Gibco, Thermo Fisher Scientific, Cat no. 31330-038), 20% KnockOut Serum Replacement (KOSR) (Gibco, Thermo Fisher Scientific, Cat no. 10828028), 1x MEM Non-Essential Amino Acids Solution (MEMNEAA) (Lonza, Cat no. 11140050), 1x GlutaMAX Supplement (Gibco, Thermo Fisher Scientific, Cat no. 35050061), 1% Penicillin-Streptomycin (Gibco, Thermo Fisher Scientific, Cat no. 15140122), 0.385 µM 2-mercaptoethanol (Gibco, Thermo Fisher Scientific, 21985023) with the addition of ROCKi (50µM) (Tocris, Cat no. 1254) for 4-6 days. Bright field images were collected during these time periods.

Brain organoids were generated with a modified version of the protocol of given in [32]. Briefly hiPSCs were plated on low-attachment U-bottom 96 well plate and cultured in ESCM supplemented with ROCKi (50µM) and 4ng/ml Human FGF-basic Recombinant Protein (Gibco, Thermo Fisher Scientific, Cat no. PHG0024). At day 6, formed embryoid bodies (EBs) were transferred into Neural Induction Media (NIM) composed of DMEM/F-12 supplemented with 1x N-2 MAX Media Supplement (R&D Systems, Cat no. AR009), 1x GlutaMAX, 1x MEMNEAA, 1 µg/ml Heparin (Sigma-Aldrich, Cat no. H3149). At day 10, developed EBs were embedded in Matrigel Basement Membrane Matrix (Corning, Cat no. 354234) cultured in Cerebral Organoid Differentiation Media (CODM) composed of 1:1 DMEM/F12 and Neurobasal Medium (Gibco, Thermo Fisher Scientific, Cat no. 21103049) supplemented with 0.5x N-2 MAX Media Supplement, 1x GlutaMAX, 0.5x MEMNEAA, 1x B-27 Supplement minus vitamin A (Gibco, Thermo Fisher Scientific, Cat no. 12587010), 2.5 µg/ml insulin (Gibco, Thermo Fisher Scientific, Cat no. I9278), 0.1925 µM 2-mercaptoethanol for 4 days. At the end of the incubation period, 3D clusters were started to incubate in CODM supplemented with B-27 Supplement with vitamin A (Gibco, Thermo Fisher Scientific, Cat no. 17504044). At this step 3D clusters were differentiated into BOs in three-different culture conditions including static culture, orbital-shaker and microfluidic chip. Dynamic conditions for orbital-shaker (Miulab GSP-20, P.R.China) were set as 85 rpm mixing and 0.35 µl/min media flow for microfluidic chip. All groups were cultured for 30 days at 37 °C incubator under 5% CO2 atmosphere and 95% humidity.

### 2.2 Fabrication of microfluidic chips

Microfluidic chips providing dynamic culture conditions for organoids by applying laminar medium flow were designed with AutoCAD (Autodesk) and CorelDRAW (Corel Co.). Chips were fabricated by stacking poly (methyl methacrylate) (PMMA) layers cut with laser cutter (Epilog-MINI) and bonded with double side adhesive (3M, 468MP, USA). Sterilization of microfluidic chips was performed by 70% ethanol and 30 minutes of UV exposure in laminar flow cabinet.

### 2.3 Immunofluorescence staining

hiPSC derived EBs and BOs were fixed with 4% paraformaldehyde (Sigma-Aldrich) for 25 min at room temperature. Brain organoids were embedded in cryomatrix (OCT, Fisher Healthcare) and cryosectioned with Leica CM1950 (Leica Biosystems, Germany). EBs and BOs were permeabilized with permeabilization buffer (0.3% Triton X-100 (neoFroxx, GmbH, Cat no. 8500) (v/v) in PBS (Gibco, Thermo Fisher Scientific, Cat no. P04-36500) for 15 min. in room temperature. Then, cells were blocked with blocking buffer (0.05% Tween 20 (Sigma-Aldrich, Cat no. P7949) (v/v), 1% Bovine Serum Albumin (BSA; Sigma-Aldrich, Cat no. A2153) (w/v) in PBS) for 1 h room temperature. Subsequently, primary antibodies (α-SMA (Cell Signaling Technology, Cat no.48938S), Nestin (Proteintech, Cat no. 19483-1-AP), Sox17 (Abcam, Cat no. Ab84990), Sox2 (Bioss, Cat no. bs-0523R), Tuj1 (R&D Systems, Cat no. MAB1195), N-cadherin (Cell Signaling Technology, Cat no. 13116T) were added according to related samples and incubated for overnight at 4 °C. Cells were stained with corresponding secondary antibodies for 2 h at room temperature. Nuclei were stained with 4,6-diamidino-2-phenylindole (0.5 μg/ml) (DAPI; Neofroxx, Cat no. 1322). Samples were visualized with fluorescence microscope (Olympus IX71).

### 2.4 Creation of datasets

Experimental protocol outlining the generation of EBs and organoids utilized in creation of datasets used in OrganoLabeling is shown in Figure 1A-B. Brightfield microscopic images with 2464 × 2056 resolution were obtained from day 1 to day 6 for EBs and from day 6 to day 44 for organoids with 10x objective using inverted microscope equipped with Zeiss Axiocam 705 color camera (Axiovert Zeiss, Germany). Prior to analyses, images were resized 256x256 for convenience. Steps followed for the labeling process of images includes contrast adjusting, K-means clustering, CLAHE, binary and Otsu thresholding methods (Fig. 1C). Details of the OrganoLabeling architecture is described in the System Architecture section.

**Figure 1.**
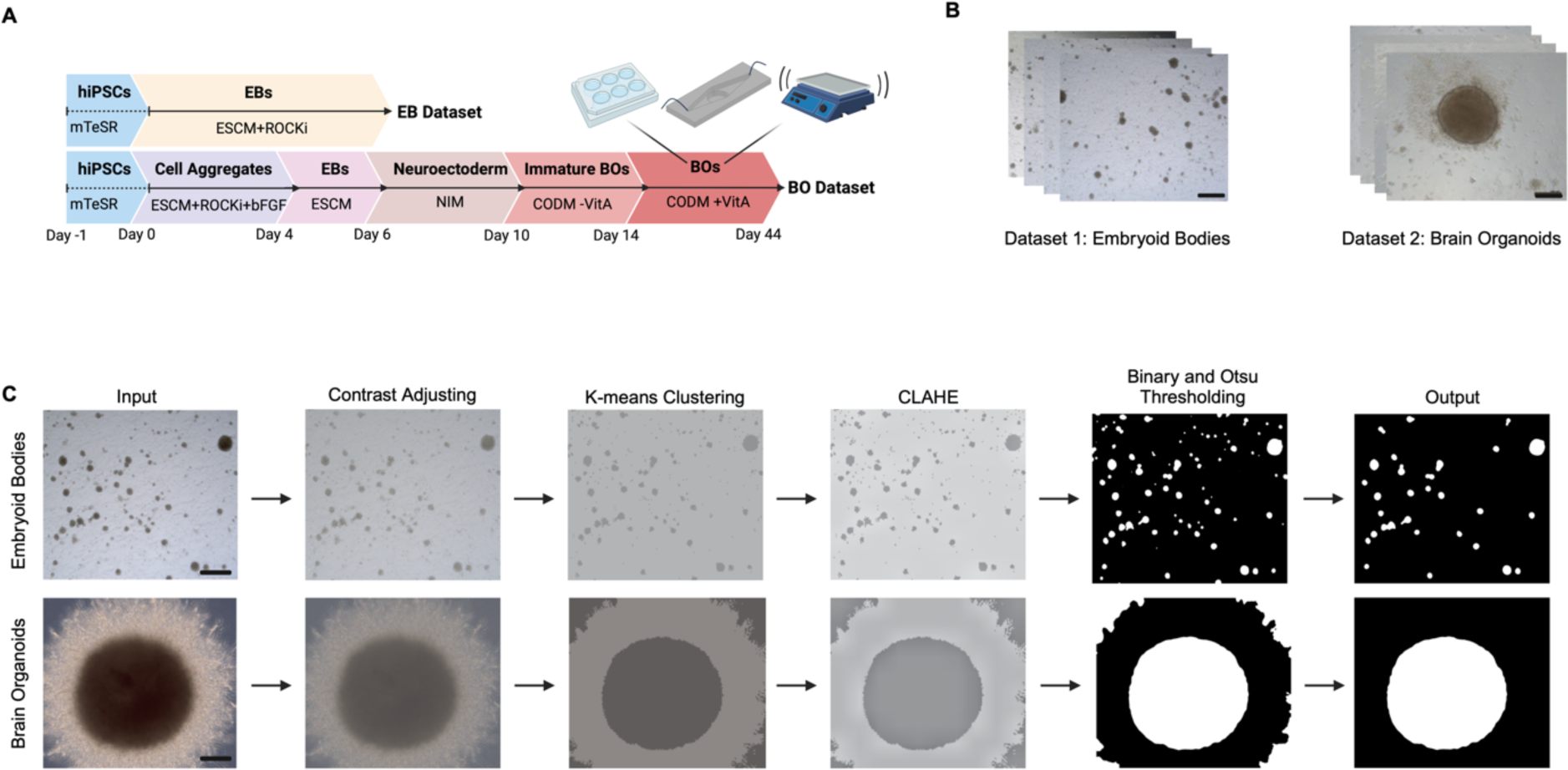
Workflow demonstrating the experimental and computational phases of OrganoLabeling A) Schematic illustration of generation of embryoid bodies and brain organoids starting from hiPSCs, B) Embryoid body and brain organoid datasets created from brightfield microscopy images (scale bars; 200µm), C) Image processing steps included in the OrganoLabeling tool.

In this study, to demonstrate the generalizability of OrganoLabeling, besides datasets created from our own experimental data we have included publicly available datasets of intestinal organoids (enteroids), generated by Beck et al. [33]. The EB dataset is formed from 165 images, BO dataset consists of 133 images and enteroid dataset consists of 299 randomly selected images (Table 2). Segmented images of the brain organoid and embryoid body dataset required for comparison with OrganoLabeling outputs were made manually using ImageJ. All datasets are publicly available on Kaggle (https://www.kaggle.com/datasets/burakkahveci/brain-organoid-and-embryoid-bodies-organolabeling).

**Table 2.** Datasets used in the OrganoLabeling system.

| Name | Image Numbers | Microscope |
| --- | --- | --- |
| Embryoid bodies | 165 | inverted |
| Brain organoids | 133 | inverted |
| Enteroids | 299 | confocal |

### 2.5 System Architecture

OrganoLabeling system architecture has been created by merging five different image processing techniques that are contrast adjustment, K-means clustering, contrast-limited adaptive histogram equalization (CLAHE), binary and Otsu thresholding methods (Figure 1C). First, input is obtained by system and contrast adjustment carried out as per entered value. Contrast adjustment plays a vital role for the separation of background and region(s) of interest (ROI). Then, OrganoLabeling system performs image segmentation with K-means clustering to be able to separate foreground object from the background. Because some images have complex background, the system also performs CLAHE. Before the thresholding step, blurring operation is performed for the better thresholding performance. Finally, ROI(s) are obtained by making binary and Otsu thresholding.

There are five parameters: contrast adjustment, blurring, clip limit and tile grid size for CLAHE, and finally the area limit. Because all these parameters can be adjusted by users, OrganoLabeling give the best output for users’ expectations.

The whole system is created by Python programming language (Version 3.9.15). Contrast of the images were adjusted with Python Image Library (Pillow). K-means clustering, CLAHE, preprocessing, and area size arranging are performed by OpenCV library. Matplotlibrary is used for visualizations of outputs. Basic mathematical operations are applied by Numpy library. U-Net models were trained with TensorFlow (2.10.0).

### 2.6 Training process with U-Net

To create the U-Net models [19], we used Keras and Tensorflow libraries. Figure 2 shows the U-Net architecture. It comprises repeated double 3x3 convolution layers, each followed by a 2x2 max pooling operation for down sampling. After each down sampling, the number of feature channels are doubled which increases up to 1024 at the end of the first half of the architecture. In the second half, each up-sampling step is followed by a 2x2 up-convolution and a concatenation with the corresponding level of the first half. Final layer uses a convolution to reduce the feature map depth to the desired number of classes.

**Figure 2.**
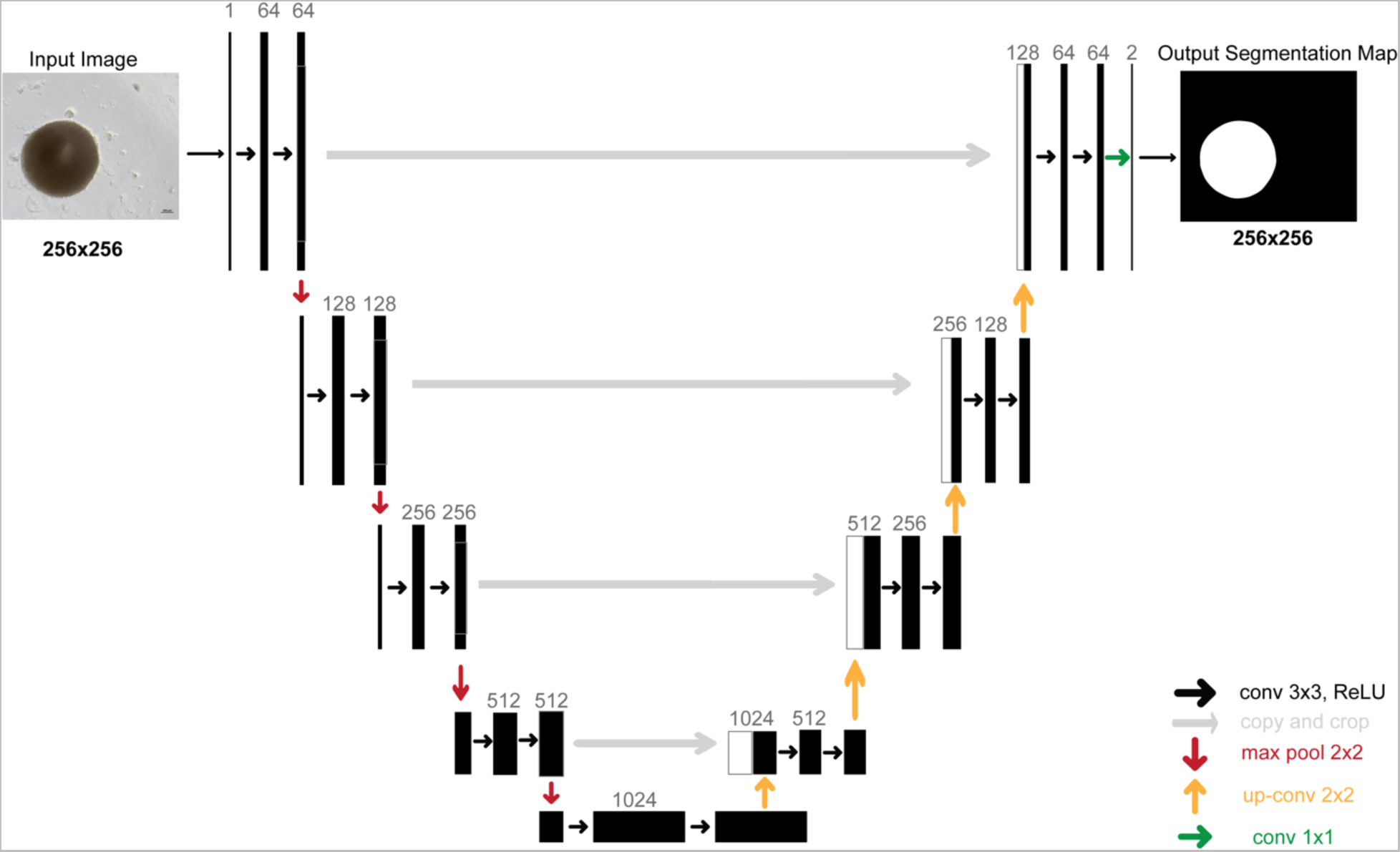
U-Net architecture.

We resized the dataset to 256x256 pixel since the U-Net model works at this resolution. In our training, we performed train and test dataset split (80/20%) and we applied 5-fold cross-validation, which corresponds to shifting the test set at each fold.

The loss function is designed as a pixel-wise softmax over the final feature map.

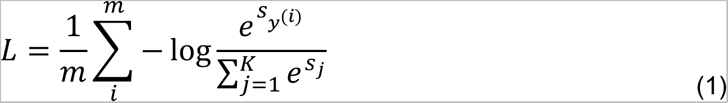

where *m* is the number of samples (pixels) and *K* is the number of classes. For pixel *i*, *y*^(*i*)^ denotes the correct class label.

We built U-Net models for three different datasets. The number of images in train and test dataset, the number of epochs, and the number of batch sizes are given in Table 3. The experiments in the study were carried out with Apple MacBook M1 Max (64 GB RAM).

**Table 3.** Dataset and model information for U-Net.

| Dataset Name | Train – Test Dataset<br>Image Numbers | Epoch | Batch Size | Loss | Optimizer | Learning<br>Rate |
| --- | --- | --- | --- | --- | --- | --- |
| Embryoid body | 131 - 34 | 100 | 8 | Binary Cross-<br>entropy | Adam | 1e-3 |
| Brain organoid | 106 - 27 | 20 | 4 | Binary Cross-<br>entropy | Adam | 1e-3 |
| Enteroid | 239 - 60 | 20 | 4 | Binary Cross-<br>entropy | Adam | 1e-3 |

### 2.7 Evaluation metrics

#### Intersection over Union (IoU)

Intersection over Union is an evaluation metric that is used for segmentation and object detection tasks. The IoU metric measures the similarity between two sets of pixels, one set being the ground truth (true segmentation mask) and the other set being the predicted result (segmentation output of the model). There are three significant terms, namely true positive (TP), false positive (FP) and false negative (FN), used to calculate IoU for the segmentation task (Table 4).

**Table 4.** Terms for calculation of IoU and DC.

|  |  |
| --- | --- |
| True Positive | Number of pixels that are correctly classified as object pixels |
| False Positive | Number of pixels that are incorrectly classified as object pixels |
| False Negative | Number of pixels that are part of the ground truth object but not predicted as being part of an object |

The formula of IoU metric is given below.

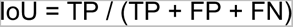

The IoU metric is a value between 0 and 1, with a value of 1 indicating a perfect segmentation match between the ground truth and predicted result.

#### Dice Coefficient (DC)

Dice Coefficient is a metric that measures the similarity or overlap between two sets. It is used to evaluate model performance in image segmentation and binary classification. Similar to IoU, its value changes between 0 and 1.

The formula of DC metric is given below.

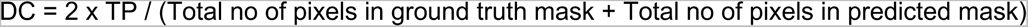

## 3. Results

We developed the OrganoLabeling tool, which quickly segments images of 3D cellular structures such as embryoid bodies and organoids. Here, we first report the characterization of EBs and BOs and further demonstrate the performance of OrganoLabeling, which has been compared to manually segmented images made by an expert researcher.

### 3.1. Characterization of embryoid bodies and brain organoids

Embryoid bodies are 3D aggregates of stem cells that are capable of generating three germ layers [34]. EBs were generated from hiPSC in 6 days of suspension culture. We confirm the differentiation capacity of hiPSC into EBs by demonstrating germ layers formed with positivity for α-SMA, Nestin and Sox17 representing the presence of mesoderm, ectoderm, and endoderm, respectively (Figure 3A). Generated EBs have diameter with range of 100-400 µm.

**Figure 3.**
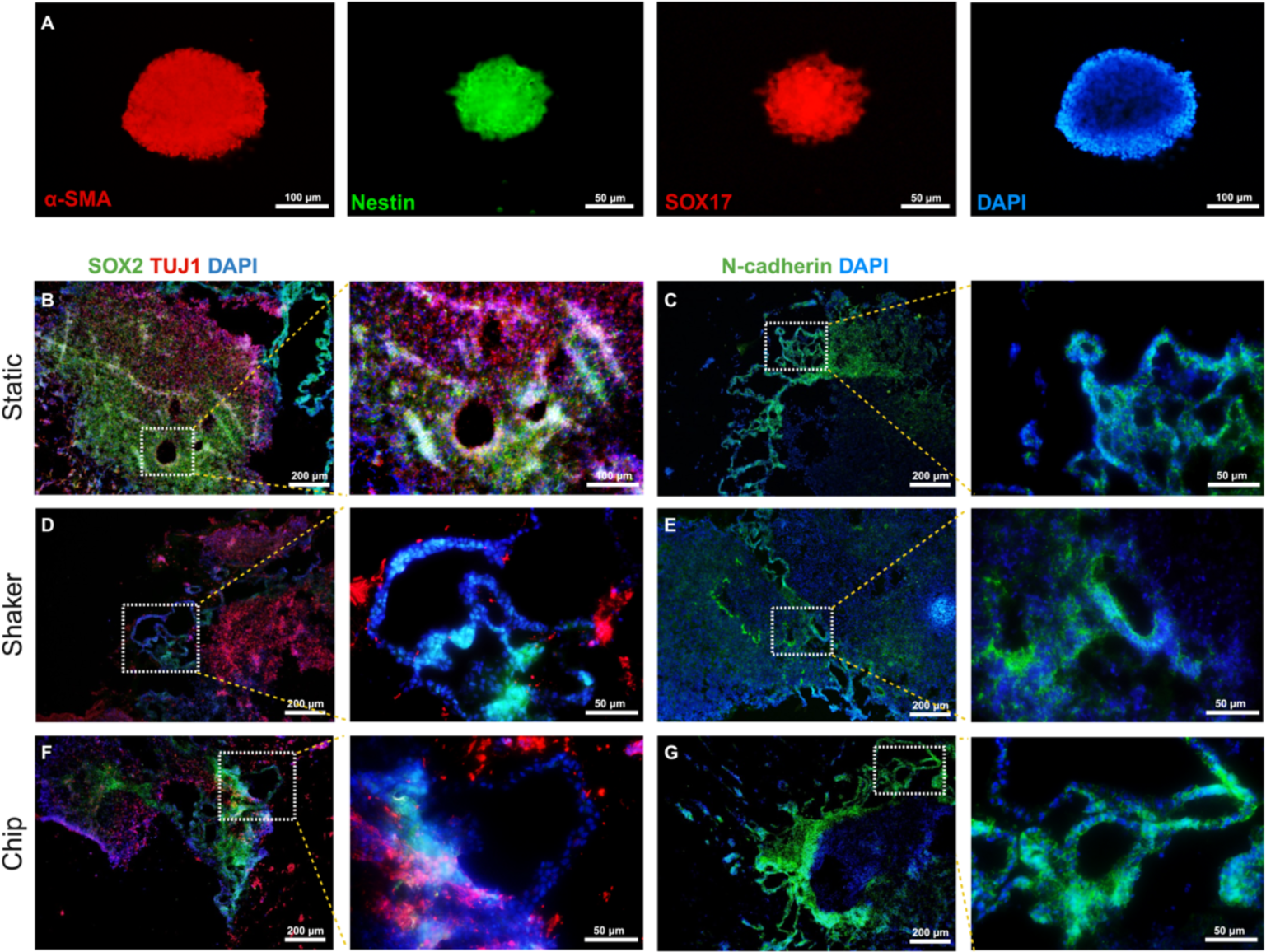
Immunochemical characterization of embryoid bodies and 30 days old brain organoids generated under three different conditions. A) Embryoid bodies stained to demonstrate the presence of three distinct germ layers; with α-SMA (mesodermal marker), Nestin (ectodermal marker), Sox17 (endodermal marker) and DAPI (cell nucleus), B) brain organoid generated under static condition stained with Sox2 and Tuj1, C) brain organoid generated under static condition stained with N-cadherin, D) brain organoid generated using orbital shaker stained with Sox2 and Tuj1, E) brain organoid generated using orbital shaker stained with N-cadherin, F) brain organoid generated using microfluidic chip stained with Sox2 and Tuj1, G) brain organoid generated using microfluidic chip stained with N-cadherin. (Right panels; magnification of dashed zones)

Brain organoids have been shown to be generated under static and dynamic (constant shaking or microfluidic chips) conditions [32, 35, 36]. To demonstrate the capacity of OrganoLabeling here we generated BOs from hiPSCs either in static culture (Fig.3 B-C), or dynamic culture conditions provided by orbital shaker (Fig. 3D-E) or microfluidic chip (Fig. 3F-G). hiPSC derived BOs were first differentiated towards the ectodermal lineage which, further give rise to neuronal tissue. The neuro-differentiation capacity of generated organoids was achieved through Sox2 expression demonstrating the presence of neural progenitors mostly observed in apical surface of organoids. Tuj1 expressing neurons originated from the neural progenitors localize separately from progenitor zone. Typical for BOs N-cadherin is expressed in the apical membrane surrounding cavities reminiscent of ventricles. Collectively this data indicates that organoids generated in different conditions have complex heterogenous regions identifying BOs.

### 3.2 Applying OrganoLabeling to different organoid datasets

Initially, we conducted OrganoLabeling on the dataset comprising EBs (Figure 4). We evaluated the outputs obtained with OrganoLabeling using expert-generated reference data and the IoU and DC parameter. Figure 4A shows examples from each evaluation metric range: 100-90%, 89-80%, and 79-50%. In Figure 4B, the percentage distributions of the outputs obtained from the OrganoLabeling application in the entire dataset is provided. Throughout the distribution obtained from the entire data set, the output metrics were concentrated in the 79-50 range in terms of IoU, while they showed a balanced distribution in terms of DC metric. Average IoU and DC were obtained from the EB dataset as 71% and 83%, respectively (Table 5).

**Figure 4.**
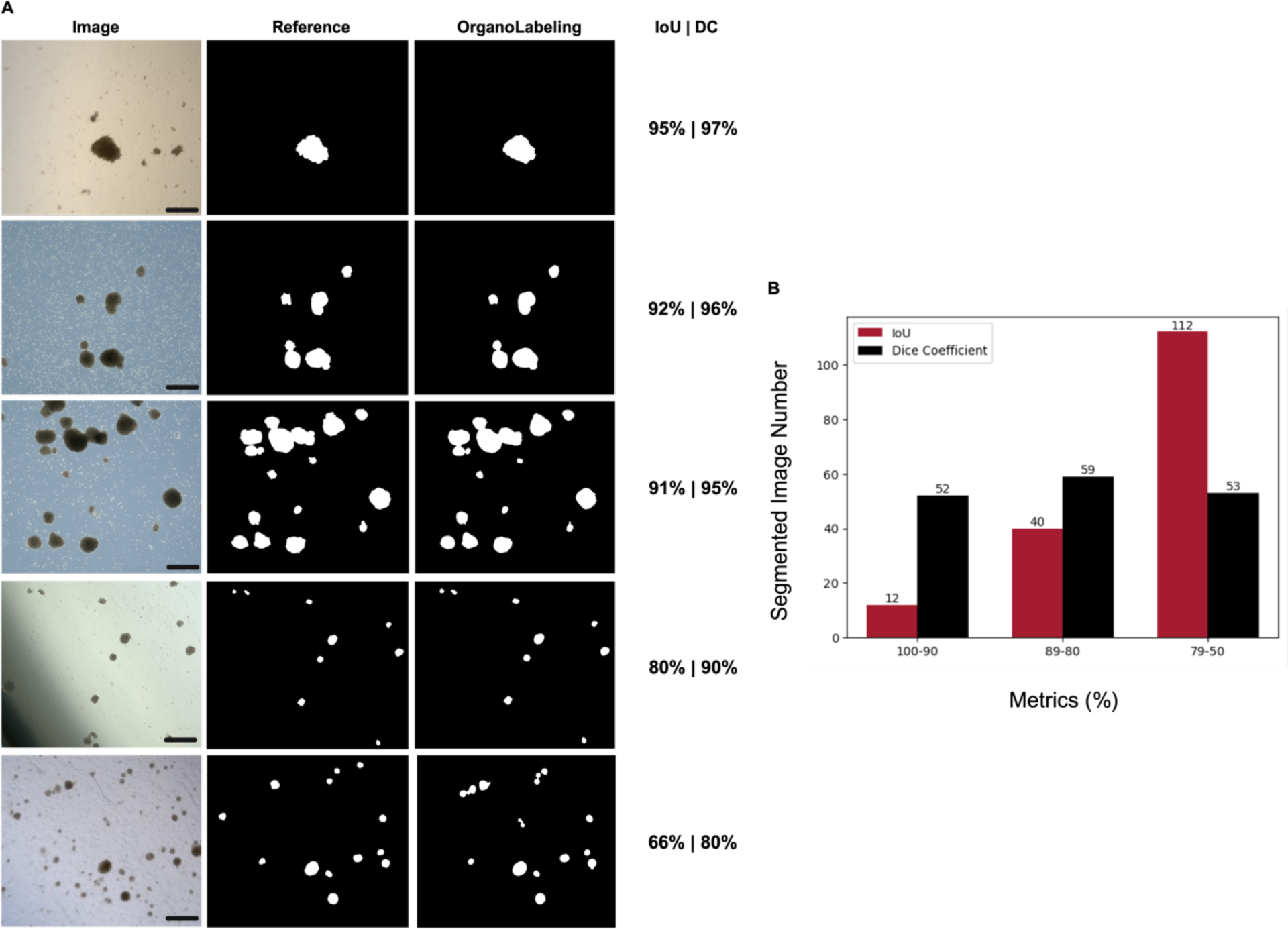
Application of OrganoLabeling to bright field images of embryoid bodies and comparison with reference segmentation. A) Examples of each evaluation metric distribution from the embryoid dataset (scale bars; 200µm). B) Metrics ranges of images segmented by OrganoLabeling compared to reference outputs.

**Table 5.** Mean IoU and DC for OrganoLabeling.

| Dataset | Dataset Size | Mean IoU (%) | Mean Dice (%) |
| --- | --- | --- | --- |
| Embryoid body | 165 | 0.71 | 0,83 |
| Brain organoid | 133 | 0.91 | 0.92 |
| Enteroid | 299 | 0.88 | 0.94 |

**Table 6.** Comparison of U-Net results of OrganoLabeling segmented and reference images.

| Datasets | U-Net Segmentation with OrganoLabeling (IoU %) | U-Net Segmentation with Reference (IoU %) | U-Net Segmentation with OrganoLabeling (DC %) | U-Net Segmentation with Reference (DC %) |
| --- | --- | --- | --- | --- |
| Brain organoid | $0.96 \pm 0.02$ | $0.94 \pm 0.03$ | $0.98 \pm 0.004$ | $0.95 \pm 0.01$ |
| Embryoid body | $0.98 \pm 0.004$ | $0.98 \pm 0.004$ | $0.98 \pm 0.08$ | $0.98 \pm 0.008$ |
| Enteroid | $0.96 \pm 0.01$ | $0.97 \pm 0.01$ | $0.96 \pm 0.01$ | $0.91 \pm 0.08$ |

OrganoLabeling was also applied to the BO dataset (Figure 5). The highest IoU and DC obtained from the outputs were determined as 98% and 97%, respectively, and the images with the five different IoU and DC percentages are given in Figure 5A. We obtained the highest number of outputs in the BO dataset in the range of 100-90. We found the output numbers in the 50% percentile to be quite low. In addition, the average IoU rate for the entire dataset was found to be 91%, while the average DC rate was found to be 92% (Table 5). This indicates that the outputs of OrganoLabeling are similar to the images created by human researchers. More importantly, the difference between the reference images and OrganoLabeling outputs does not affect the performance of deep learning models as will be seen in Section 3.3. This proves OrganoLabeling outputs can be used to train deep segmentation models which eliminates the need for manually labeled images.

**Figure 5.**
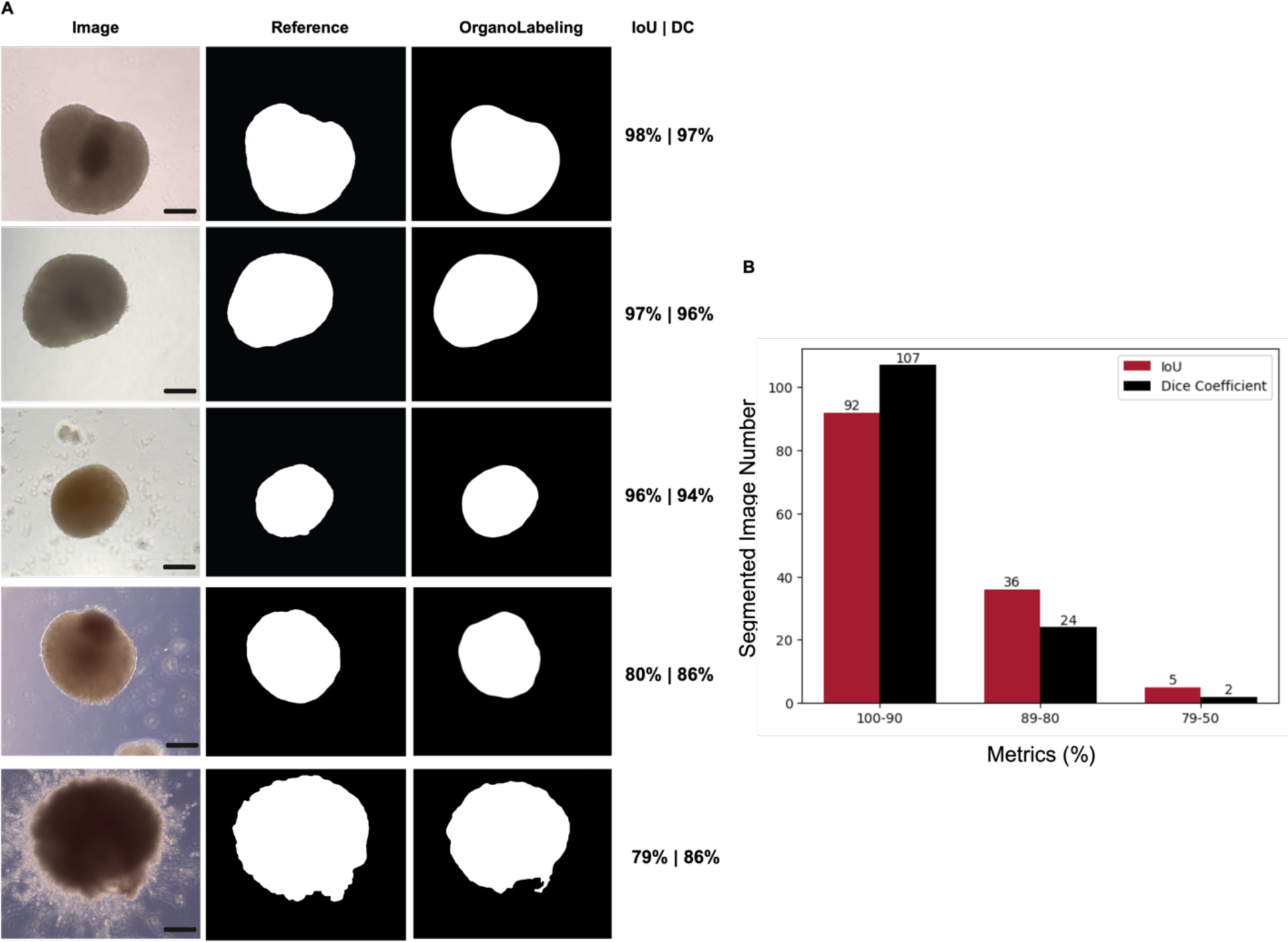
Application of OrganoLabeling to brain organoids and comparison with expert segmentation. A) Five images with the different IoU and DC in the brain organoid dataset (scale bars; 200µm). B) IoU and DC ranges of brain organoid images segmented by OrganoLabeling compared to reference outputs.

To demonstrate the generalizability of OrganoLabeling, we used the publicly available entoroid dataset [33] (Figure 6). Here OrganoLabeling showed high performance, similar to in-house generated datasets. Five examples with different percentage distribution of IoU and DC are shown in Figure 6A. OrganoLabeling reached the highest percentage in this dataset with an IoU percentage of 98%. The images in the image groups randomly taken from the public dataset and the average IoU and DC percentages obtained for each group are given in Figure 6B. When segmented images obtained with OrganoLabeling compared to images created by human researchers, we achieved an average IoU rate of 88% and DC rate of 94%. As shown in Figure 6C, the output from OrganoLabeling is predominantly in the 100-90 range. The part where the outputs are least intense is the 79-50 range.

**Figure 6.**
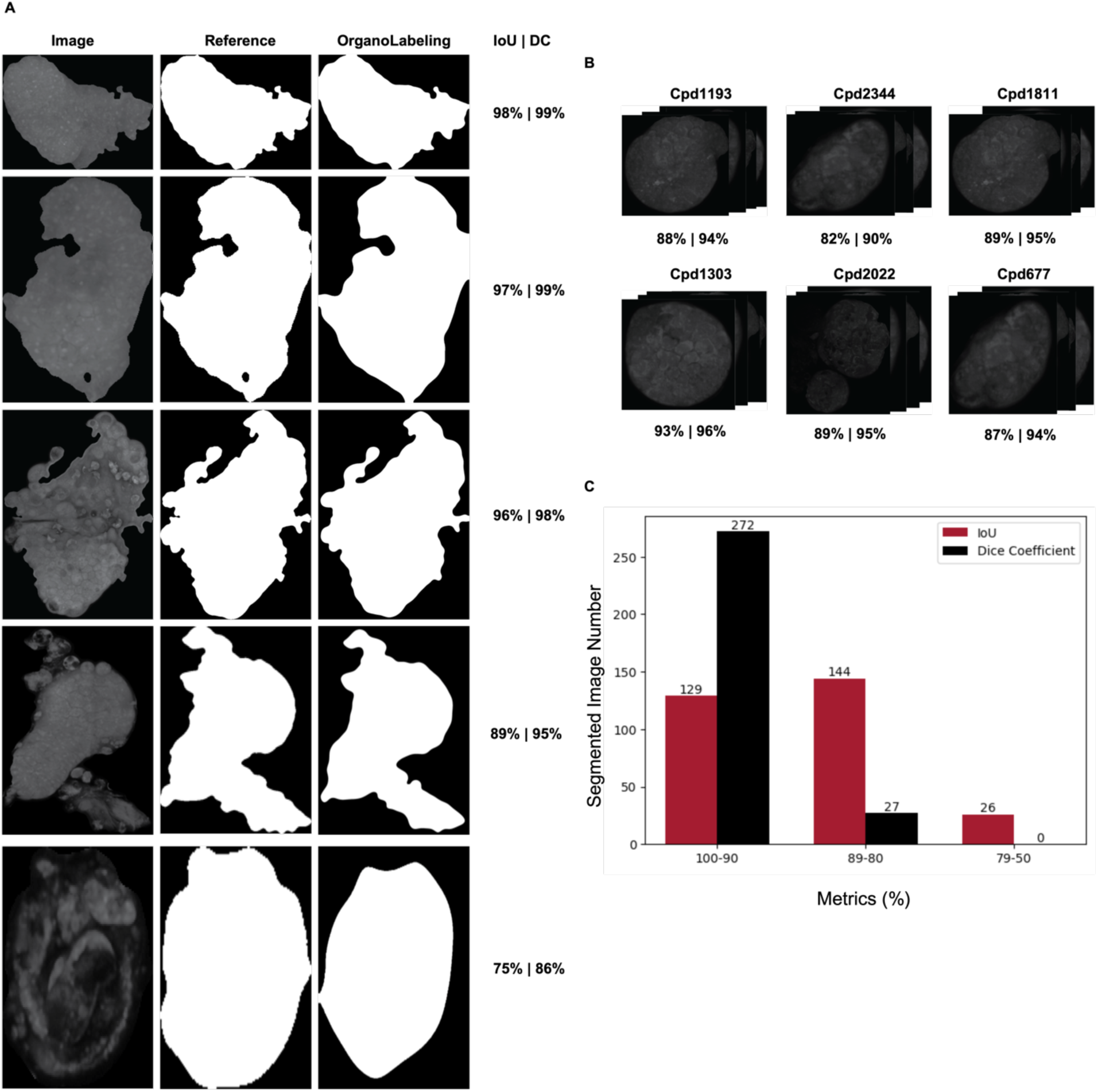
Application of OrganoLabeling to enteroid dataset and comparison with reference segmentation. A) Five images with different distribution percentage of IoU and DC in the enteroid dataset. B) Randomly selected data groups from enteroid dataset. C) IoU and DC ranges of enteroid images segmented by OrganoLabeling compared to reference outputs.

### 3.3 Comparison of U-Net model trained using OrganoLabeling segmented images with the U-Net model trained using manually segmented images

For three different datasets, we trained and tested U-Net models using OrganoLabeling segmented images and manually segmented images (Figure 7). As shown in Figure 7A, two separate U-Net models were trained and tested on the EB dataset. One model is trained with images segmented by OrganoLabeling and the other model is trained with manually segmented (reference) images. Both U-Net models achieved a 98% IoU and DC rates, indicating that OrganoLabeling can replace the manual labeling process of EB dataset. Likewise, two separate U-Net models were trained for the BO dataset. Figure 7B shows three randomly selected test images and their segmentation made by i) U-Net model trained with OrganoLabeling images and ii) U-Net trained with manually labeled images. The U-Net model trained using images segmented with OrganoLabeling achieved an IoU rate of 96% and DC rate of 98%, while the U-Net model trained using manually labeled images achieved a 94% IoU rate and 95% DC rate. Lastly, figure 7C shows the comparison for enteroid images, where U-Net models trained with OrganoLabeling segmented and manually segmented images obtained IoU rates of 96% and 97% respectively. Additionally, DC rate of OrganoLabeling segmented images is 96%, while DC rate of manually segmented images is 91%. Results indicate that U-Nets trained with OrganoLabeling output images perform at least as well as those trained with reference datasets. OrganoLabeling can be used to create datasets for training deep learning models (i.e. no need for manual pixel labeling).

**Figure 7.**
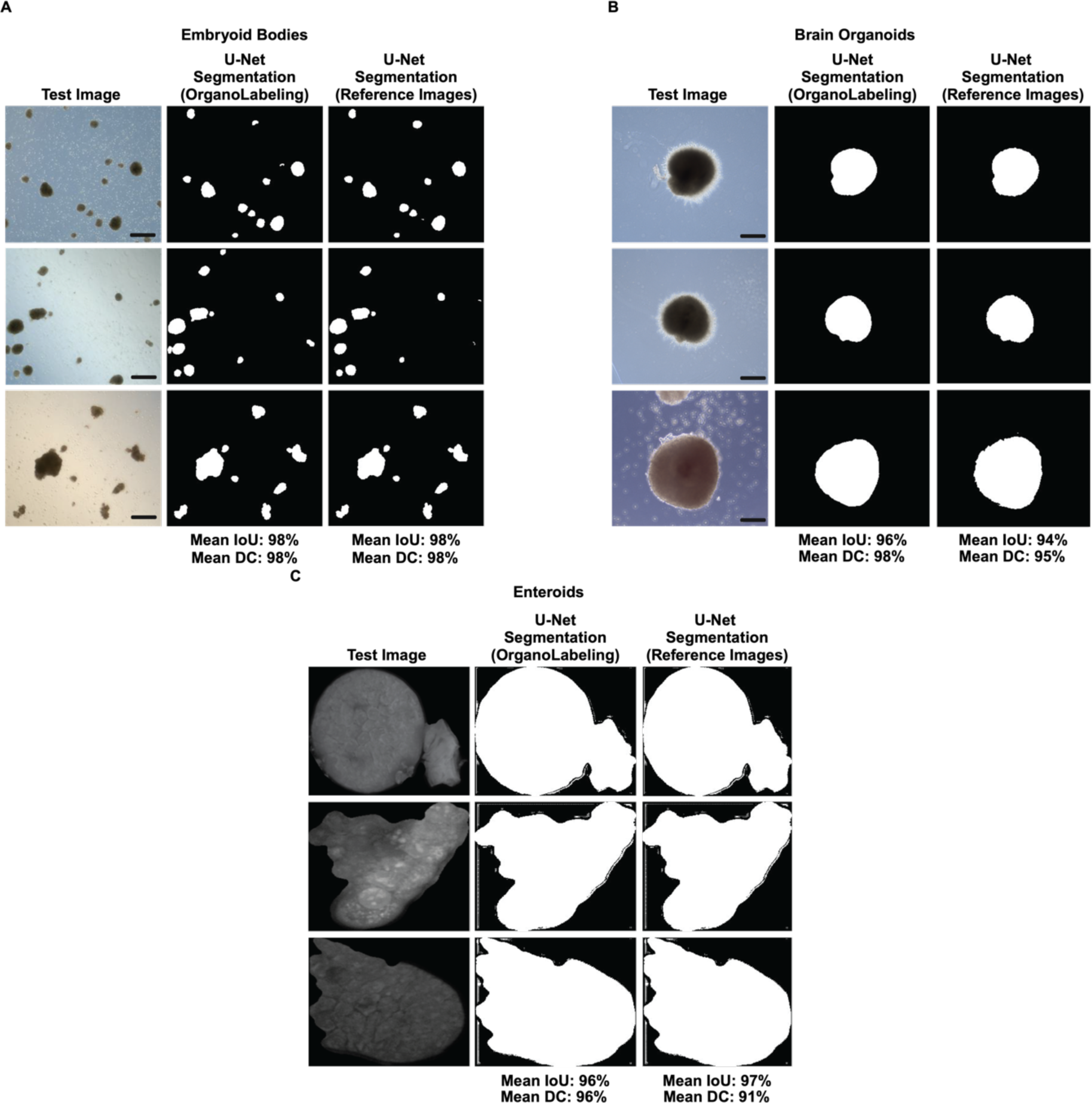
Training the U-Net model with both the dataset created by OrganoLabeling and the reference dataset and comparing the results. A) The average IoU and DC values of two different U-Net models trained for embryoid bodies and three randomly selected images and their segmented versions are given (scale bars; 200µm). B) Two different U-Net models trained for brain organoids, average IoU and DC values are shown (scale bars; 200µm). C) Two different U-Net models trained and tested for the enteroid dataset and their average IoU and DC values are given. For the training of each U-Net model, the K-Fold value was determined as 5. Comparative results of three randomly selected images have shown.

## 4. Discussion

Organoids are 3D tissues that recapitulate structural and functional features of organs and can be widely utilized in life sciences and health applications such as disease modeling, drug screening, personalized medicine and developmental biology [1]. Similar to other tissue organoids, brain organoids derived from hiPSCs have 3D morphologies formed in complex cellular composition. Starting from cell aggregates differentiation process towards brain organoids takes weeks even months with costly laborious tasks. Although applying identical differentiation protocols, the yield of generation brain organoids with distinct histological characterization may differ, indicating not all cell aggregates succeed towards the targeted organoid. Utilization of image based analytical tools may assist to select morphologically potent 3D cell clusters at early phase, eliminating number of samples thereby minimizing expenses.

Here we report an automated image labeling tool named as OrganoLabeling, which can be utilized further in segmentation of datasets used in artificial intelligence based predictive models. Such labeling tools are essential to digitalize wet lab generated biological results into processible dataset in image based computational analyses. In this study we first generated embryoid bodies that are precursors of organoids and further obtained brain organoids as relevant 3D biological samples to be employed in labeling of image datasets.

Embryoid bodies are 3D cellular structures formed during the generation of brain organoids. Their cellular composition initiates the further differentiation towards specific type of tissue, considering the germ layer origin. We have accordingly characterized the hiPSCs derived EBs for mesoderm, ectoderm and endoderm with positivity to (-SMA, Nestin and SOX17 respectively. Brain organoids were generated under static and dynamic settings using three different culture conditions as previously reported [32, 36–38]. We demonstrate the successful differentiation of hiPSCs into BO where organoids from all conditions show positivity for neuroprogenitor SOX2 and neuronal marker TUJ1 through immunohistological characterization (Figure 3). Zonal cavities representing the ventricle structures present in BOs have been identified with N-Cadherin staining. Independent from culture conditions we obtained BOs to be used as model organoids for labeling.

Although there are several labeling tools for deep learning models, these tools are both time-consuming and prone to human-based errors because they require manual labeling [39–41]. However, OrganoLabeling differs from these as a system that automates image labeling process. Morphological tracking of organoids is of great importance in the process of their creation. Deep learning-based approaches can be used for this tracking. Studies using DL models for the analysis of images obtained from organoids have been increasing in recent years [23–28]. Additionally, tools for morphological analysis of organoids have been developed using deep learning models [29, 30]. Convolutional neural networks were used for the segmentation of organoids in the tool developed by Matthews et al. They trained their system with a pancreatic organoid dataset [29].. Segmentation was carried out with the U-Net model in OrgaExtractor developed by Park et al [30]. Colon organoid was used as dataset in the study. However, training of these models is laborious and challenging due to the fact that the segmented images have to be prepared manually. In our study, we developed a no-code automated tool called OrganoLabeling.

To test our tool, we used three different datasets and trained U-Net models using the resulting segmented images. We also trained U-Net models using manually labeled images and compared the results. The U-Net models trained with segmented images generated with OrganoLabeling mostly outperformed the models trained with manually segmented images.

Two of the datasets used included mature BOs from EBs summarizing the organoid development process, while the other was a dataset generated with enteroid images. The embryoid body dataset contains 165 images, the BO dataset contains 133 images and the enteroid dataset contains 299 images. Thus, OrganoLabeling was tested with different organoid types and mostly exceeded the success of manually labelled data. Moreover, it accelerated the labeling process considerably.

OrganoLabeling has a few limitations. In the segmentation of small EBs in the EB dataset, one of the three datasets used, our system had difficulty in segmenting appropriately. In addition, there were some images taken with inverted microscopes of the EBs in the images taken from the boundary parts of the EBs where the EBs were located, which tended to be incorrectly segmented due to the shadow created by the wells. In addition, in a small number of images containing this shadow, the EBs coming into the shadow part could not be segmented by the system.

## 5. Conclusion

In this study, we developed OrganoLabeling, a tool that can generate segmented organoid images to be used for DL models for the images of cell and cellular structures. We compared the performance of OrganoLabeling with images manually segmented by experts. OrganoLabeling outputs are similar enough to the manually segmented images so that they can be used to train deep segmentation convolutional neural networks. We proved that by training a state of the art segmentation CNN (U-Net). U-Net models trained with segmented images generated with OrganoLabeling mostly outperformed the models trained with manually segmented images. Being user-friendly, OrganoLabeling is available free of charge, with user-adjustable parameters. This makes it easy to create fast and human error-free datasets for segmentation models to be trained with organoid images.

## AUTHOR CONTRIBUTIONS

B.K. and S.G. established the hypothesis and study design. E.P. and B.K. generated brain organoids and embryoid bodies and created datasets. B.K. created OrganoLabeling and carried out the computational experiments. B.K. and Y.B. trained U-Net models. All authors contributed the manuscript writing. S.G. and Y.B. edited the manuscript. All the authors read and approved the manuscript.

## ACKNOWLEDGMENTS

This work is supported by Dokuz Eylül University ADEP TSA 2023-3026 project. E.P. is fellow of YÖK 100/2000, TÜBİTAK 2211A and 2250 scholarship programs. B.K. is fellow of TÜBİTAK 2250 scholarship program.

## DATA AVAILABILITY STATEMENT

All codes and datasets are available to Kaggle (https://www.kaggle.com/datasets/burakkahveci/brain-organoid-and-embryoid-bodies-organolabeling).

